# The Juicebox Assembly Tools module facilitates *de novo* assembly of mammalian genomes with chromosome-length scaffolds for under $1000

**DOI:** 10.1101/254797

**Authors:** Olga Dudchenko, Muhammad S. Shamim, Sanjit S. Batra, Neva C. Durand, Nathaniel T. Musial, Ragib Mostofa, Melanie Pham, Brian Glenn St Hilaire, Weijie Yao, Elena Stamenova, Marie Hoeger, Sarah K. Nyquist, Valeriya Korchina, Kelcie Pletch, Joseph P. Flanagan, Ania Tomaszewicz, Denise McAloose, Cynthia Pérez Estrada, Ben J. Novak, Arina D. Omer, Erez Lieberman Aiden

## Abstract

Hi-C contact maps are valuable for genome assembly (Lieberman-Aiden, van Berkum et al. 2009; Burton et al. 2013; Dudchenko et al. 2017). Recently, we developed Juicebox, a system for the visual exploration of Hi-C data (Durand, Robinson et al. 2016), and 3D-DNA, an automated pipeline for using Hi-C data to assemble genomes (Dudchenko et al. 2017). Here, we introduce “Assembly Tools,” a new module for Juicebox, which provides a point-and-click interface for using Hi-C heatmaps to identify and correct errors in a genome assembly. Together, 3D-DNA and the Juicebox Assembly Tools greatly reduce the cost of accurately assembling complex eukaryotic genomes. To illustrate, we generated *de novo* assemblies with chromosome-length scaffolds for three mammals: the wombat, *Vombatus ursinus* (3.3Gb), the Virginia opossum, *Didelphis virginiana* (3.3Gb), and the raccoon, *Procyon lotor* (2.5Gb). The only inputs for each assembly were Illumina reads from a short insert DNA-Seq library (300 million Illumina reads, maximum length 2x150 bases) and an *in situ* Hi-C library (100 million Illumina reads, maximum read length 2x150 bases), which cost <$1000.

---

An accurate genome sequence is an essential basis for the study of any organism. To assemble a genome, a large number of DNA sequences derived from the organism of interest are overlapped with one another to create contiguous sequences, known as “contigs.” Next, linking information – derived from a wide variety of sources, such as mate-pairs, physical maps, and read-clouds – is used to order and orient these contigs into “scaffolds.” Errors often arise throughout the assembly process. Contigs may mistakenly concatenate two sequences (a ‘misjoin’). Scaffolds can contain errors in contig order (a ‘translocation’) or orientation (an ‘inversion’). Examples of such errors can be found in the best available reference genomes for many species (Robert B. Norgren 2013; Shearer et al. 2014; Tang et al. 2014; Chen et al. 2015; Davey et al. 2016; Utsunomiya et al. 2016; Schneider et al. 2017; Korlach et al. 2017). Consequently, inexpensive methods for identifying and correcting assembly errors are crucial for the generation of accurate assemblies (Salzberg and Yorke 2005; Phillippy, Schatz, and Pop 2008; Gnerre et al. 2009; Tsai, Otto, and Berriman 2010; Salzberg et al. 2012; Hunt et al. 2013; Gurevich et al. 2013; Bradnam et al. 2013; Simão et al. 2015; Fierst 2015; Muggli et al. 2015; Yuan et al. 2017; Harewood et al. 2017). Of course, improved error correction procedures can also reduce the amount of input data required, and thereby the cost of genome assembly.

Hi-C, a method for determining the 3D configuration of chromatin, is emerging as a valuable source of data for genome assembly (Lieberman-Aiden et al. 2009; Burton et al. 2013; Session et al. 2016; Peichel et al. 2016; Bickhart et al. 2017; Dudchenko et al. 2017; Mascher et al. 2017). When visualized, Hi-C data is typically represented as a heatmap. This heatmap is generated by partitioning a reference genome assembly into loci of fixed size; each heatmap entry indicates the frequency of contact between a pair of loci. When a chromosome is correctly assembled, sequences that are adjacent in the assembly are also in close physical proximity, leading to the appearance of a bright band of elevated contact frequency along the diagonal of the Hi-C heatmap. Conversely, when there are errors in a reference assembly, they are often visually obvious as anomalous patterns in the heatmap (Rao, Huntley et al. 2014; Harewood et al. 2017; Dudchenko et al. 2017; Lapp et al. 2017). Thus, in addition to its use as an input to automated assemblers, Hi-C can also facilitate the visual identification of errors in a genome assembly.

Recently, we introduced Juicebox, a set of tools that facilitate the visual exploration of Hi-C heatmaps across a wide range of scales (Durand, Robinson et al. 2016). Here, we introduce “Assembly Tools” (see Fig. 1), a new module in the Juicebox desktop application that extends the Juicebox interface in order to facilitate interactive genome assembly and reassembly using Hi-C data. Assembly Tools enables users to superimpose the positions of contigs or scaffolds in a reference assembly on top of the Hi-C heatmap, making assembly errors easier to find. When assembly errors are found, users can correct them, using a simple point-and-click interface, in a matter of seconds. Both the heatmap and the reference genome are updated in real-time to reflect these changes. Using Assembly Tools, users can improve genomes and reduce the cost of genome assembly.

**Figure 1:**
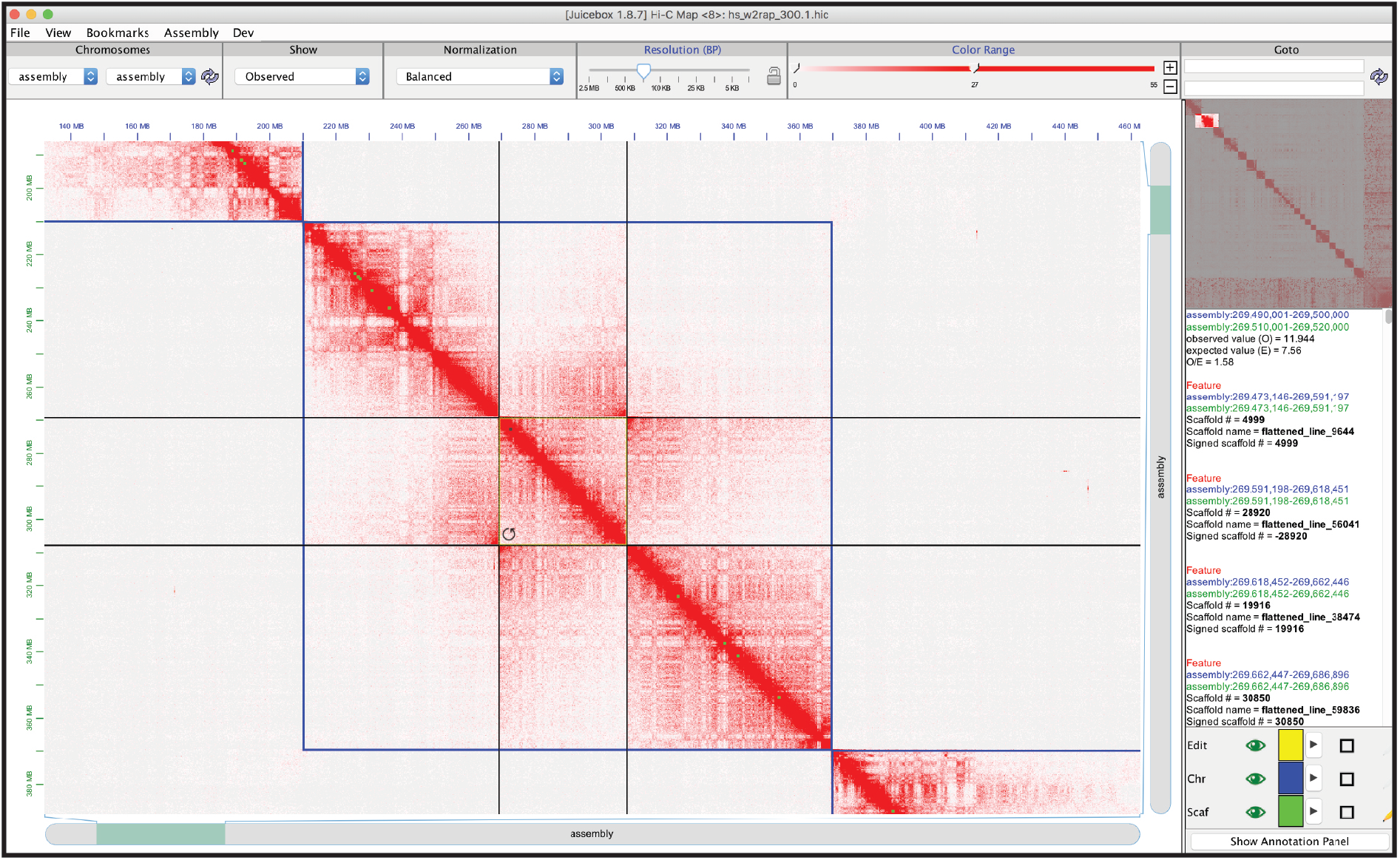
Juicebox Assembly Tools module enables visualization and interactive manipulation of genome assemblies. A screenshot of Juicebox Assembly Tools module zoomed in to 100-kb resolution on a region of a human genome assembly. A mouse prompt for inversion is shown appearing in the lower left of the selected genomic interval. While using the Assembly Tools, users can continue to employ standard Juicebox functions. The toolbar at the top allows users to quickly navigate between different views, normalizations, and resolutions as well as to load and save assembly files. At the top right, a mini-map shows the whole chromosome at low resolution. Below, hover text shows data for scaffolds in the selected genomic interval (yellow and black highlight). Two-dimensional features representing scaffold and chromosome boundaries are superimposed on the main map. Their appearance can be modified using Annotation panel in the lower right.

To begin, a user needs to specify an assembly to be modified. Like the 3D-DNA algorithm, Assembly Tools uses a custom format, *.assembly*, that can be quickly generated from a *.fasta* file by an accompanying command-line tool. The user also needs relevant Hi-C data in the *.hic* format. In practice, this will often entail performing a Hi-C experiment in the organism of interest (Rao, Huntley et al. 2014), generating between 0.01X and 20X coverage, and running Juicer (Durand, Shamim et al. 2016).

Once the assembly and Hi-C dataset have been loaded via a pull-down menu, the user can begin to identify and correct errors. For instance, a translocation typically manifests as an extremely bright bowtie motif pointing horizontally or vertically, whose midpoint corresponds to two loci that are proximate in the genome but lie far apart in the assembly (Rao, Huntley et al. 2014; Dudchenko et al. 2017; Harewood et al. 2017). By clicking-and-dragging, a user can highlight the desired genomic interval, and – with a single click – move the interval to the desired position in the assembly (see Fig. 2a). Similarly, an inversion error – when the sequence of bases in a genomic interval is reversed – often manifests as a bowtie parallel to the diagonal (Rao, Huntley et al. 2014; Dudchenko et al. 2017). By clicking at the center of this motif, users can invert the selected interval (see Fig. 2a). Finally, a misjoin typically manifests as a point along the diagonal of the Hi-C heatmap where the upper-right and lower-left quadrants are extremely depleted, reflecting the lack of physical proximity between the erroneously concatenated loci (Dudchenko et al. 2017). Such errors can be resolved by selecting the affected scaffold and clicking on the position of the misjoin. The scaffold is then split in two in the reference genome assembly, allowing the two resulting scaffolds to be separately manipulated until they are correctly placed (see Fig. 2a). In addition, a third, short scaffold, containing the misjoined sequence itself, is excised and relocated to the end of the reference genome assembly, where anomalous scaffolds are kept for future reference. The boundaries of superscaffolds, such as chromosomes or chromosome arms, can be indicated by clicking between two scaffolds when no interval is currently selected. To simplify the above correction process, the mouse prompt changes to indicate the operation that is possible at any given moment: a circular arrow for inversion; a straight arrow for translocation; scissors for misjoin excision; and an angle to introduce a superscaffold boundary (see Fig. 2a).

**Figure 2:**
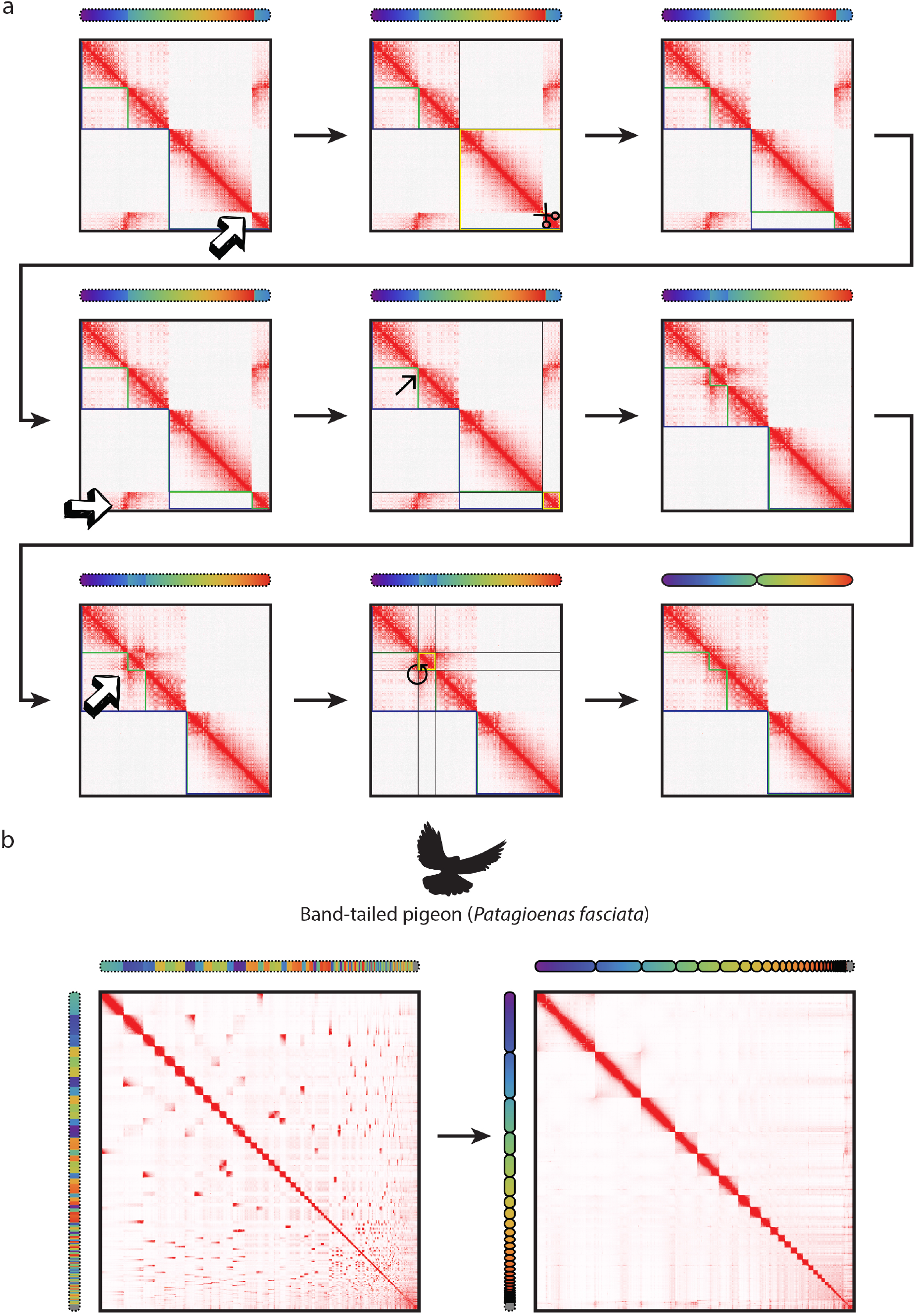
Hi-C maps allow for visual identification and interactive correction of errors in genome assemblies. (a) Here we show a contact matrix generated by aligning a GM12878 Hi-C data set (HIC001 library from (Rao, Huntley et al. 2014)) to a simulated genome assembly containing several errors. Each ‘pixel’ in the map indicates the frequency of contact between a pair of loci in the assembly. As the original assembly is being modified interactively using Juicebox Assembly Tools, the changes are reflected in the heatmap. This process continues until no more anomalous signals can be found on the heatmap. The simulated assembly was created by deliberately introducing errors into the sequence of two chromosome-length scaffolds from hg19 (chromosomes 2 and 4). The position of the loci according to hg19 is shown using chromograms. For the purpose of illustration, gaps have been removed from hg19 sequence. Anomalies in the Hi-C heatmap associated with 3 types of misassemblies (misjoin, translocation and inversion) are indicated by hand-drawn errors on the left-side panels. The interaction with Juicebox Assembly Tools (cut, paste and invert) and the accompanying mouse prompt are indicated in the middle panels. The resulting heatmaps are shown on the right-side panel. The simulated assembly consists of two chromosomes (boundaries outlined with blue annotations). The first chromosome comprises two pieces (boundaries outlined with green), while the second one comprises one. (b) Interactive genome assembly using Juicebox Assembly Tools results in chromosome-length scaffolds for the band-tailed pigeon. The left-side panel shows the draft genome assembly generated by aligning a HiC data set to the draft genome assembly GCA_002029285.1 (Murray et al. 2017), which was used as input into Assembly Tools. The right-side panel illustrates the contact map for resulting assembly. Corresponding loci are indicated using chromograms.

After each change to the reference genome assembly, the Hi-C heatmap that is being displayed by Juicebox is updated accordingly. Crucially, the Juicebox Assembly Tools module does not recalculate the *.hic* file storing the Hi-C heatmap at each step, a process which could take many hours (Durand, Shamim et al. 2016). Instead, the new heatmap can be thought of as a rearrangement of the pixels in the old heatmap, permuting its rows and columns. The Assembly Tools module tracks this permutation, updating it each time a change is made to the reference genome assembly.

While using the Assembly Tools module, users can also continue to employ standard Juicebox functions; for instance, they can modify the color scale, or zoom in and out (Durand, Robinson et al. 2016). A user can save the current state of the genome assembly as a new *.assembly* file. When the user is finished, a simple script can be run from the command line in order to apply this assembly file to the original reference assembly *.fasta* file, producing a corresponding assembly sequence.

To illustrate the use of the Assembly Tools module, we re-examined data from a very recent study which assembled the genome of the band-tail pigeon *(Patagioenas fasciata)*, the closest living relative of the extinct passenger pigeon (Murray et al. 2017). This assembly incorporated Illumina and *in vitro* Hi-C (Chicago), yielding a scaffold N50 of 20 Mb. We generated *in situ* Hi-C data for the band-tailed pigeon (239M read pairs, 66X coverage). We then ordered the extant scaffolds from largest to smallest, loaded them into Juicebox, and performed an interactive genome assembly, resulting in chromosome-length scaffolds (scaffold N50: 76 Mb). (See Figs. 2b and S1.)

Although the Juicebox Assembly Tools module can be used independently, it can also be used as a validation and refinement system for the output of our automated 3D-DNA pipeline, which uses Hi-C data to improve genome assemblies. By adding a manual validation and refinement step, reliable genome assemblies can often be generated using less input data, reducing the cost of *de novo* genome assembly.

To illustrate the use of 3D-DNA and Juicebox Assembly Tools in tandem, we developed a procedure for assembling mammalian genomes with chromosome-length scaffolds for under $1000 (see Fig. 3a). Our procedure involves three steps, and can be performed by a single person in roughly 10 days. First, we generate a PCR-free short insert DNA-Seq library, and sequence 300 million paired-end Illumina reads (2x150 bases). This corresponds to roughly 30X coverage for a typical (3Gb) mammal. These reads are assembled into a draft assembly using the software package w2rap (B. Clavijo et al. 2017; B. J. Clavijo et al. 2017). Second, we generate an *in situ* Hi-C library (Rao, Huntley et al. 2014), and sequence 100 million paired-end Illumina reads, corresponding to roughly 10X coverage for a typical mammal, which are used to improve the draft assembly by providing both as inputs to 3D-DNA (Dudchenko et al. 2017). Finally, we validate and refine the improved assembly using Juicebox Assembly Tools. Note that this procedure does not require advance knowledge of the exact size of the mammalian genome, or of the number of chromosomes.

**Figure 3:**
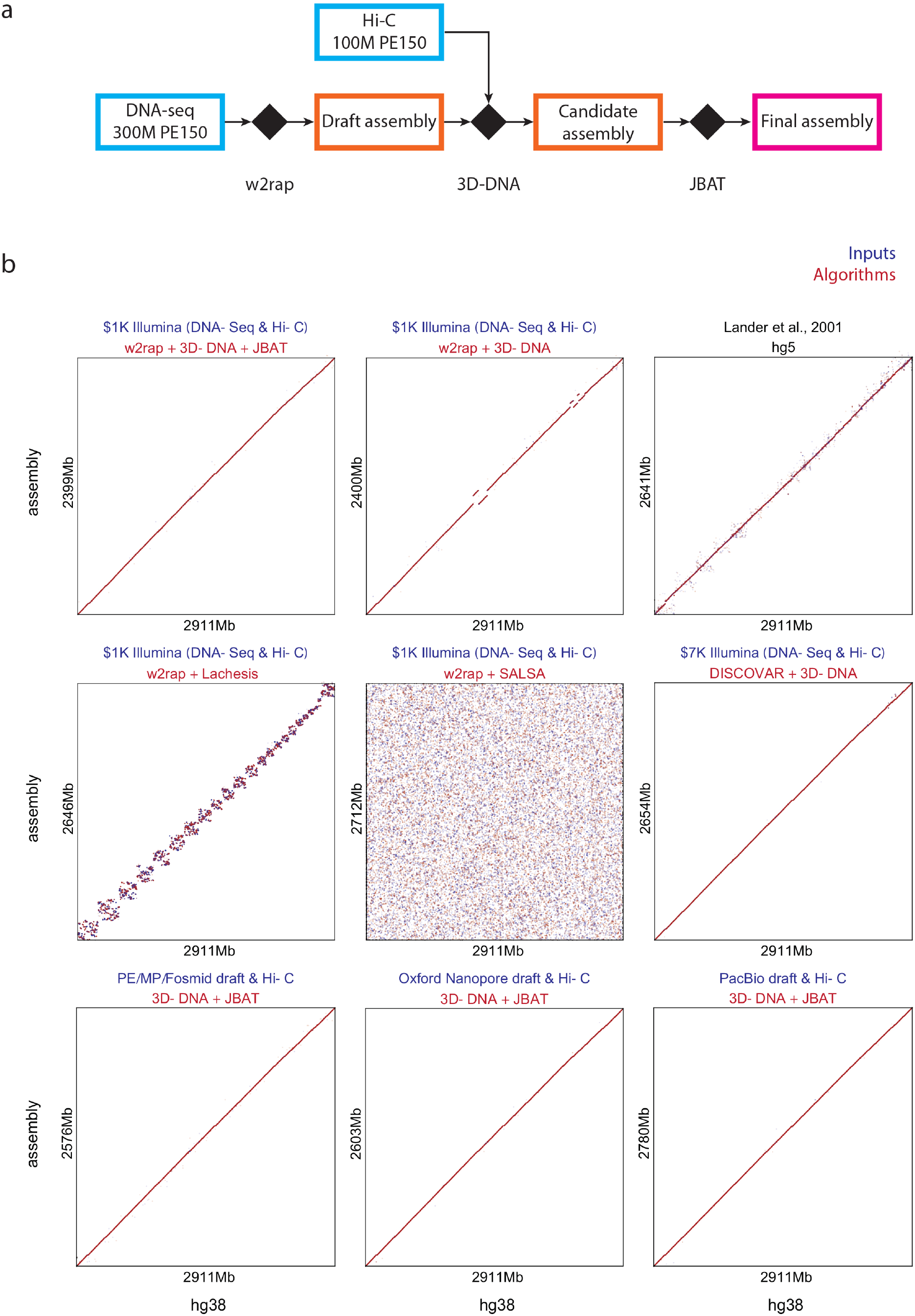
*De novo* assembly of mammalian genomes with chromosome-length scaffolds for $1000. (a) A schematic representation of a $1000 mammalian sequencing procedure. First, we generate a PCR-free short insert DNA-Seq library, and sequence 300 million paired-end reads (2x150). These reads are assembled into a draft assembly using the software package w2rap (B. Clavijo et al. 2017; B. J. Clavijo et al. 2017). Second, we generate an *in situ* Hi-C library (Rao, Huntley et al. 2014), and sequence 100 million paired-end reads (2x150). Note that the Hi-C library can often be sequenced with shorter reads. When using PE150 all necessary data, in principle, can be obtained from a single lane of an Illumina HiSeq X instrument. The data are used to improve the draft assembly using 3D-DNA (Dudchenko et al. 2017). Finally, we validate and refine the improved assembly using Juicebox Assembly Tools (JBAT). (b) Dotplots showing alignment of several different human genome assemblies to hg38 chromosome-length scaffolds, genome-wide view. The hg38 reference (NCBI accession number GCA_000001405.23) is shown on the X-axis. Each dot represents the position of a 1kb sequence chunk aligned to hg38. The dotplots are subsampled such that every 50^th^ chunk is displayed. The color of the dots reflects the orientation of individual alignments with respect to hg38 (red indicates a match, whereas blue indicates disagreement). Alignment was performed using BWA (Li and Durbin 2009). The assemblies shown are, left to right and top to bottom: (1) hs-1k genome assembly presented in this study; (2) 3D-DNA assembly algorithm (Dudchenko et al. 2017) applied to $1000 data, without Juicebox Assembly Tools review; (3) hg5 genome assembly produced in 2001 by the International Human Genome Consortium (Lander et al. 2001); (4) Lachesis algorithm for Hi-C scaffolding (Burton et al. 2013) applied to $1000 data; (5) SALSA algorithm for scaffolding long-read assemblies with Hi-C (Ghurye et al. 2017) applied to $1000 data; (6) Hs2-HiC genome assembly, produced with PE250 short insert DNA-Seq data and Hi-C, scaffolded with 3D-DNA, see (Dudchenko et al. 2017); (7-9) 3D-DNA with Juicebox Assembly Tools procedure applied to more expensive DNA-Seq input data types: a collection of Illumina libraries with varying insert sizes including paired-end, mate-pair and fosmid libraries (Gnerre et al. 2011); Oxford Nanopore reads (Jain et al. 2017) and Pacific Biosciences long reads. Note that the automatic 3D-DNA chromosome splitter failed to split the assembly at the coverage associated with hs-1k input. As such we have split the 3D-DNA output into 23 chromosomes manually. The order and orientation of scaffolds inside chromosomes was not changed. All assemblies except for hg5 are NA12878 genome assemblies that have been scaffolded using the same Hi-C data: the first 100 million reads from the HIC001 library, whose generation and initial analysis was reported in (Rao, Huntley et al. 2014).

To confirm the accuracy of the resulting genomes, we used our procedure to generate a *de novo* assembly of a human genome, see Fig. 3b and S2. We took 300 million raw reads from the NA12878 dataset shared by the Genome in a Bottle Consortium (NIST NA12878 HG001 HiSeq 300x) and added 100 million reads from the GM12878 Hi-C library published in (Rao, Huntley et al. 2014, HIC001). The resulting assembly, hs-1k, contains 23 chromosome-length scaffolds which together span 88,735 contigs (contig N50: 36,914) and 2,399,853,403 sequenced bases, comprising 85.2% of the genome assembly. It also contains 1,169 small scaffolds, spanning 1,652 contigs (contig N50: 21,506) and 397,814,093 bases, comprising the remaining 14.1% of the assembly. These small scaffolds contain contigs that could not be positioned reliably using Hi-C data, typically because they were very short.

Comparison of hs-1k with the human genome reference, hg38, showed that the 23 chromosome-length scaffolds in hs-1k correctly corresponded to the 23 human chromosomes. Of the 37,074 scaffolds that were incorporated into chromosome-length scaffolds in hs-1k and that could be uniquely placed in hg38, 99.97% (comprising 99.99% of the sequenced bases) were assigned to the correct chromosome. Together, the chromosome-length scaffolds in hs-1k spanned 99.34% of the length and 82.43% of the sequence in the chromosome-length scaffolds of hg38.

Next we examined the accuracy of the ordering of these chromosome-length scaffolds. When pairs of draft scaffolds assigned to the same chromosome were examined, the order in hs-1k matched the order in hg38 in 99.86% of cases. For scaffolds that were adjacent in hs-1k, the order matched hg38 96.03% of the time, reflecting the fact that Hi-C data is less effective at determining fine structure order; when the two scaffolds were longer than 100kb, the rate increased to 99%. Similarly, the orientation of scaffolds in hs-1k matched the orientation in hg38 91.64% of the time, with the errors again arising mostly from short scaffolds.

It is interesting to compare hs-1k to the draft genome reported by the International Human Genome Sequencing Consortium (hg5) (Lander et al. 2001). The hs-1k genome assembly contains ∼10% less sequence in chromosome-length scaffolds then hg5. By contrast, the fine structure order is considerably more accurate in hs-1k. For instance, 23.1% of 1-kilobase intervals are in the wrong orientation in hg5; for hs-1k, the value is 5.2%, a 4.5-fold decrease. Note, however, that hg5 was a draft genome, and subsequent finishing steps on each chromosome greatly improved its fine structure accuracy. (See Fig. 3b.)

It is also interesting to examine the effect of replacing 3D-DNA in the above assembly strategy with two other automated Hi-C-based assembly algorithms, Lachesis (Burton et al. 2013) and SALSA (Ghurye et al. 2017). When provided the same inputs that were used for hs-1k, SALSA, which is designed to work with long-read assemblies, did not meaningfully improve upon the input. Lachesis successfully anchored many of the contigs but did not provide an accurate chromosome-scale ordering. Consequently, subsequent refinement with Juicebox Assembly Tools proved unrealistic in both cases. (See Fig. 3b.)

Having validated the $1000 genome assembly procedure, we implemented it in order to generate *de novo* assemblies of three mammals for which no assembly has been published to date: the common wombat, *Vombatus ursinus*, the Virginia opossum, *Didelphis virginiana*, and the common raccoon, *Procyon lotor* (see Fig. 4a). In each case, the result was a set of chromosome length scaffolds: for wombat, the procedure generated 7 chromosome-length scaffolds (vu-1k), spanning 83.7% of the sequenced bases, with a total contig length of 2.74Gb; for Virginia opossum, the procedure generated 11 chromosome-length scaffolds (dv-1k), spanning 79.9% of the sequenced bases, with a total contig length of 2.67Gb; and for raccoon, 19 chromosome-length scaffolds (pl-1k), spanning 77.6% of sequenced bases, with a total contig length of 1.94Gb. The new assemblies facilitate the study of karyotype evolution in marsupials and carnivores (see Figs. 4b, S3 and Supplementary table S2).

**Figure 4:**
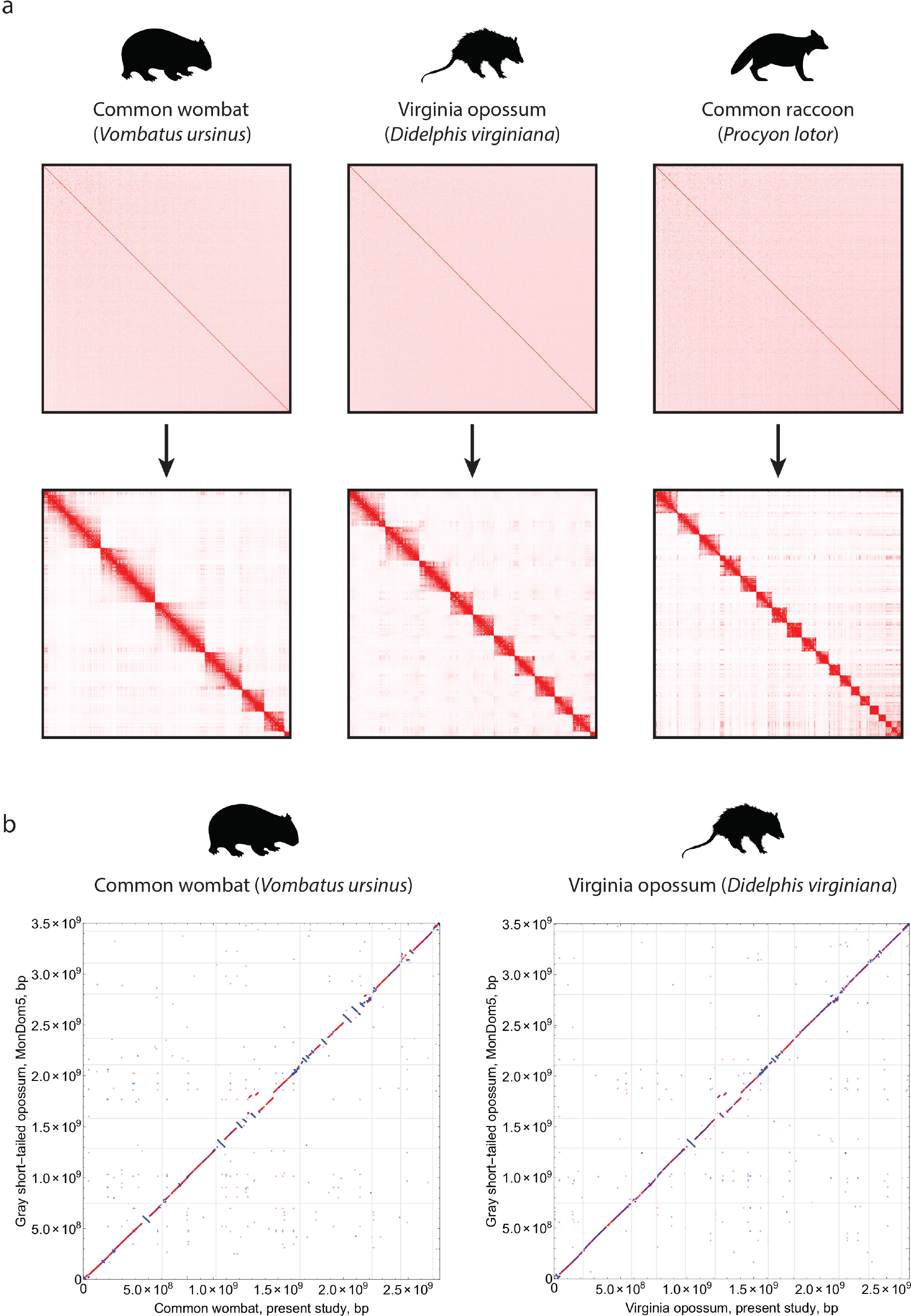
*De novo* genome assembly of three mammalian species – common wombat, Virginia opossum, and common raccoon – each using 400 million Illumina reads (2x150). (a) Draft assemblies are visualized on the top panel while final, chromosome-length assemblies are shown in the bottom. Only scaffolds larger than 15kb are displayed. Both the draft and the final assemblies are visualized using the Hi-C data employed for 3D-DNA genome assembly. (b) Chromosome-length *de novo* assemblies for the common wombat and Virginia opossum facilitate the analysis of karyotype evolution in marsupials. For this analysis, the gray-tailed opossum genome assembly (GCF_000002295.2) and the common wombat (vu-1k) and Virginia opossum (dv-1k) *de novo* assemblies were aligned using the LastZ alignment algorithm (Robert S. Harris 2007) using “--notransition --step=20 -nogapped” command options; the gray-tailed opossum genome assembly was used as a target. Here, we show alignment blocks with scores larger than 50,000 for the common wombat and larger than 65,000 for the Virginia opossum (Robert S. Harris 2007), with direct synteny blocks colored red, and inverted blocks colored blue. Chromosome order and orientation has been modified in order to facilitate the comparison.

Taken together, these findings demonstrate that the procedure we describe can be reliably employed in order to generate *de novo* assemblies of mammalian genomes with chromosome-length scaffolds.

Strikingly, the cost of the *de novo* genome assembly strategy described above is comparable to the present cost of human genome resequencing, in which short insert size DNA-Seq reads from an individual are compared to the existing human reference genome. To achieve human genome resequencing for $1000, Illumina introduced a strategy that generates up to 400 million paired-end DNA-Seq reads (2x150 bases) on a HiSeq X instrument (llumina, Inc. 2016). The *de novo* genome assembly strategy described above uses extremely similar inputs, simply replacing 100 million of the 400 million paired-end DNA-Seq reads with *in situ* Hi-C reads.

Of course, the genome assemblies generated using the strategy we describe can be further improved. For example, the genomes are not “finished” (Consortium 2004): it would be valuable to incorporate additional sequence into the chromosome-length scaffolds to fill gaps, and to correct errors in the fine-scale ordering of small adjacent contigs. Finally, although we did not encounter this issue with the hs-1k assembly, it can sometimes be difficult to correctly orient genomic intervals separated by extremely large gaps, such as chromosome arms separated by very large centromeres. These issues can be partially alleviated by additional short read Illumina data. For instance, doubling the number of PE150 reads included in our marsupial assemblies led to a larger number of sequenced bases in chromosome length scaffolds (common wombat, vu-2k: 2.72Gb→2.87Gb; Virginia opossum, dv-2k: 2.67Gb→2.85Gb). More expensive data types, such as long-read DNA sequences, can be employed to further improve the genome assembly (see Fig. 3b). The above methods are compatible with all data types of which we are aware, and we provide examples for a variety of such use cases in Table 1 and Table S3.

**Table 1:**
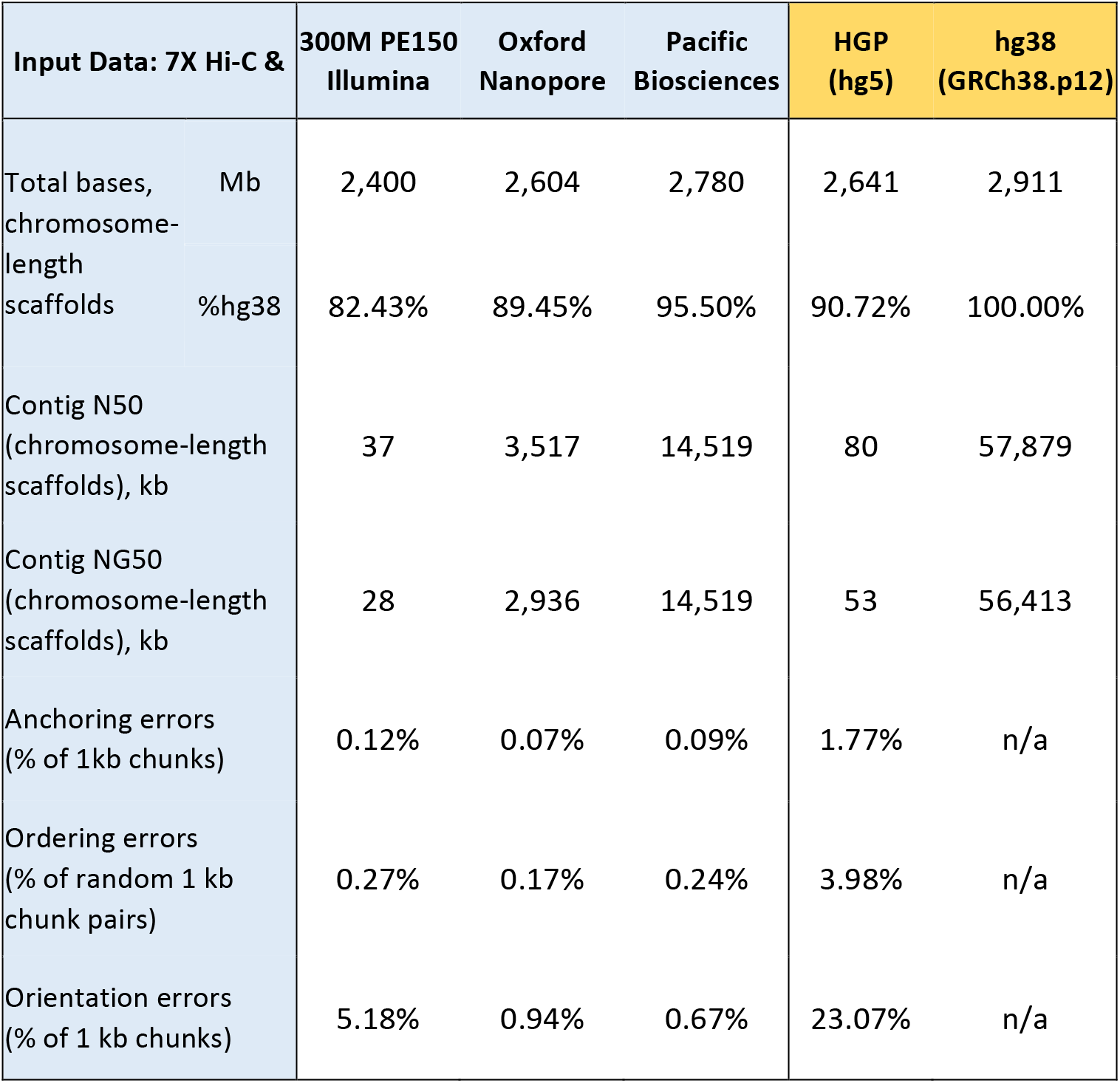
The results of 3D-DNA and Juicebox Assembly Tools using various input data types as compared to the reference genomes produced by the International Human Genome Consortium in 2001 (hg5) and the most recent reference hg38 (GRCh38.p12). The Oxford Nanopore NA12878 assembly is based on draft shared by (Jain et al. 2017); the Pacific Biosciences NA12878 assembly is based on a high-quality contigs GCA_002077035.2 shared by the McDonnell Genome Institute. Assumed genome size for NG50 estimates is 3031.04 Mb. See also Supplementary table S3.

Finally, we note that this methodology is not restricted to mammals, and can be applied successfully to many other clades. Depending on the size of the genome of interest, more or less input data may be required. Similarly, the approach could be used to generate personalized genomes in a clinical setting.

**SOFTWARE AVAILABILITY AND DOCUMENTATION OF TOOL REVIEW**. The Assembly Tools module is available as a part of the Juicebox data visualization system for Hi-C, which can be downloaded at aidenlab.org/juicebox. The code, which is available at https://github.com/theaidenlab/juicebox is open source, and is licensed under the MIT license. Genomes, datasets, tutorials and other procedures associated with this publication are available at aidenlab.org/assembly.

## ACKNOWLEDGMENTS

We acknowledge the McDonnell Genome Institute at the Washington University School of Medicine for sharing the NA12878_prelim.2.1 contigs, which were used for one of the assemblies in Table 1. We thank André Soares for providing the data from the UCSC Paleogenomics Lab on behalf of the Murray et al. group. We acknowledge David Oehler and Jean Paré of the Wildlife Conservation Society and Andrea Lee, Jess Jimerson, Katie Plaeger and Erin Neer of the Houston Zoo for their help with sample collection. We thank Christine Molter, Maryanne Tocidlowski, Lauren Howard and Judilee Marrow for veterinary work with the mammalian subjects. We also thank Chad Nusbaum and Andreas Gnirke for comments on the manuscript, and David Weisz, Alyssa Blackburn, Sheikh Russell for computational assistance. Finally, we thank Terry Leatherland, Grace Liu, Loic Fura and Victoria Nwobodo for access to a high RAM IBM E880 server. This work was supported by a Center for Theoretical Biological Physics postdoctoral fellowship to O.D., an NIH New Innovator Award (1DP2OD008540-01), an NSF Physics Frontiers Center Award (PHY-1427654, Center for Theoretical Biological Physics), the Welch Foundation (Q-1866), an NVIDIA Research Center Award, an IBM University Challenge Award, a Google Research Award, a Cancer Prevention Research Institute of Texas Scholar Award (R1304), a McNair Medical Institute Scholar Award, an NIH 4D Nucleome Grant U01HL130010, an NIH Encyclopedia of DNA Elements (ENCODE) Mapping Center Award UM1HG009375, the President's Early Career Award in Science and Engineering to E.L.A.

---

**Supplementary table S1:** Material for the mammalian assemblies described in this study. The material has been collected in the Houston Zoo by performing three opportunistic blood draws, one draw per each mammal, secondary to veterinary and/or husbandry procedures scheduled to maintain health and welfare of the animals. Each blood draw (∼1ml) was split in two to prepare DNA-Seq and Hi-C libraries. For the DNA-Seq library, we extracted DNA using QIAGEN DNeasy Blood & Tissue Kit, following the manufacturer's protocols. The DNA was sheared and prepared for Illumina sequencing using the TruSeq DNA PCR-Free kit, following the manufacturer's protocols. For Hi-C, peripheral blood mononuclear cells were separated using a Percoll gradient. The cells were crosslinked, and *in situ* Hi-C libraries were prepared in accordance with (Rao, Huntley et al. 2014). The band-tailed pigeon sample (frozen liver) was provided by the Wildlife Conservation Society. Tissue was crosslinked and dounce homogenized. Nuclei were purified on a sucrose gradient and processed to prepare *in situ* Hi-C libraries as previously described (Rao, Huntley et al. 2014). *This is the isolate that was used to generate the Hi-C data. The draft assembly has been created from a different individual (Murray et al. 2017).

|  | Common wombat | Virginia opossum | Common raccoon | Band-tailed pigeon |
| --- | --- | --- | --- | --- |
| **Binomial:** | *Vombatus ursinus* | *Didelphis virginiana* | *Procyon lotor* | *Patagioenas fasciata* |
| **Isolate:** | Lilly | Luna | Gracie | N2016-0142* |
| **Sex:** | Female | Female | Female | Male |
| **Source:** | Houston Zoo | Houston Zoo | Houston Zoo | WCS |
| **Tissue type:** | Blood | Blood | Blood | Liver |
| **Amount:** | 2ml | 0.75ml | 0.75ml | 10mg |
| **Notes:** | - | leucistic, wild caught | - | hatchling |

**Supplementary table S2:** Assembly statistics for *de novo* mammalian genomes produced in the current study.

|  | **vu-1k** | **dv-1k** | **pl-1k** |
| --- | --- | --- | --- |
| **Binomial:** | *Vombatus ursinus* | *Didelphis virginiana* | *Procyon lotor* |
| **Total bases:** | 3,273,318,797 | 3,339,913,245 | 2,505,094,826 |
| **Total bases in chr-length scaffolds:** | 2,739,633,857 | 2,669,132,684 | 1,944,533,151 |
| **Total bases in chr-length scaffolds (% total):** | 83.70% | 79.92% | 77.62% |
| **Contig N50, bp:** | 53,404 | 29,622 | 34,230 |
| **Scaffold N50, bp:** | 557,133,179 | 227,394,598 | 114,539,748 |

**Supplementary table S3:**
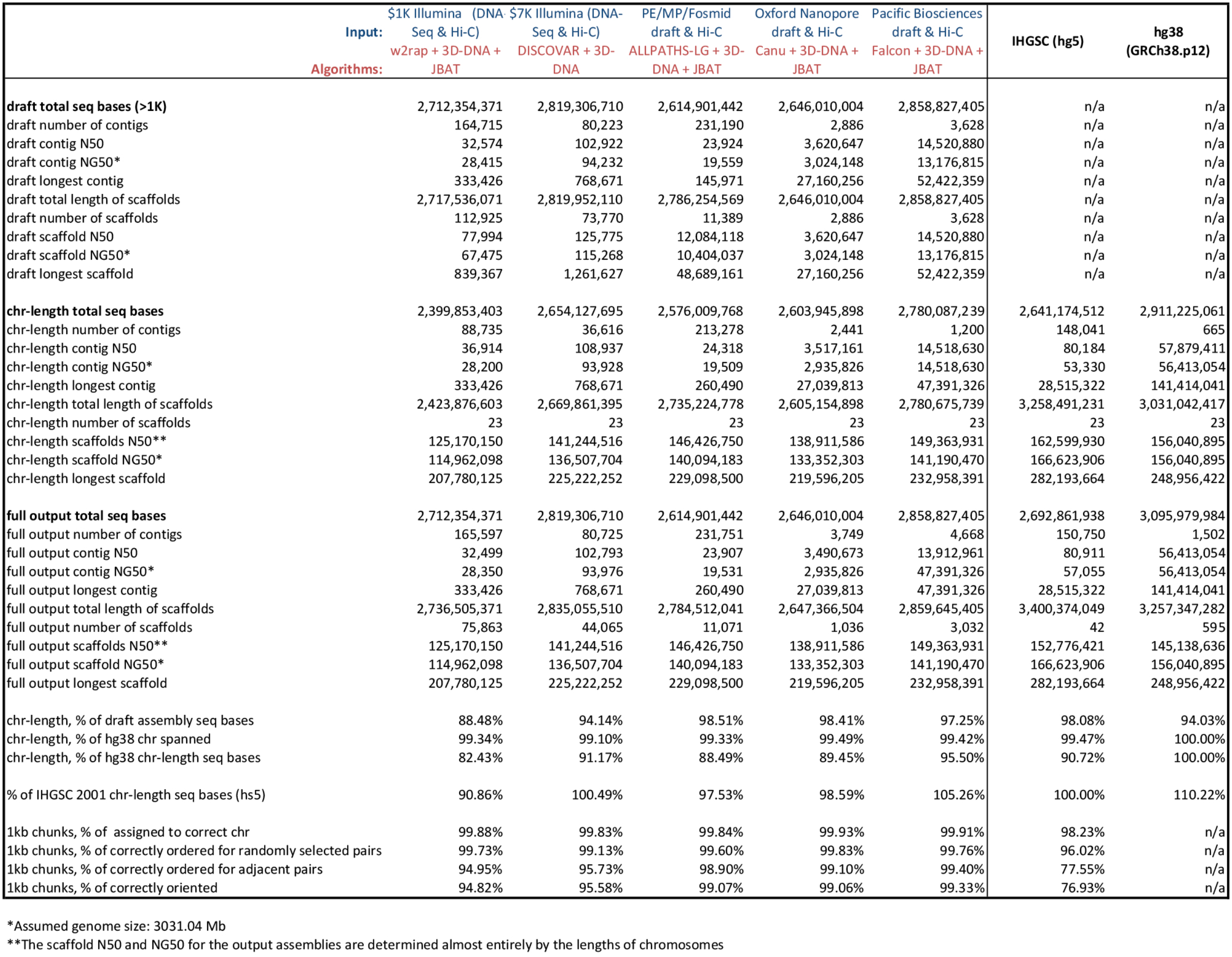
Detailed statistics for human genomes assembled with 3D-DNA and Juicebox Assembly Tools as compared to several reference genomes. Here we present some more detail on the assemblies shared in Table 1 and add a comparison to two more NA12878 genome assemblies: one generated using short insert size Illumina PE250 data and assembled using DISCOVAR *de novo* (Weisenfeld et al. 2014; Love et al. 2016) and 3D-DNA (Dudchenko et al. 2017); and another one generated with 3D-DNA and Juicebox Assembly Tools (JBAT) from a collection of short read Illumina libraries with varying insert sizes from (Gnerre et al. 2011). Accuracy statistics listed include: (1) the percentage of 1kb sequences that are placed in chromosome-length scaffolds and corresponds to the “correct” chromosome (identified by whole-genome alignment) in hg38; (2) the percentage of randomly selected pairs of 1kb sequences assigned to the same chromosome-length scaffold in the assembly that are ordered in agreement with hg38; (3) the percentage of consecutive pairs of 1kb sequences that are ordered in agreement with hg38; (4) the percentage of 1kb sequences that are oriented in agreement with hg38. Only sequences uniquely aligning to hg38 (mapq>=60) are considered in all of the analyses.

**Figure S1:**
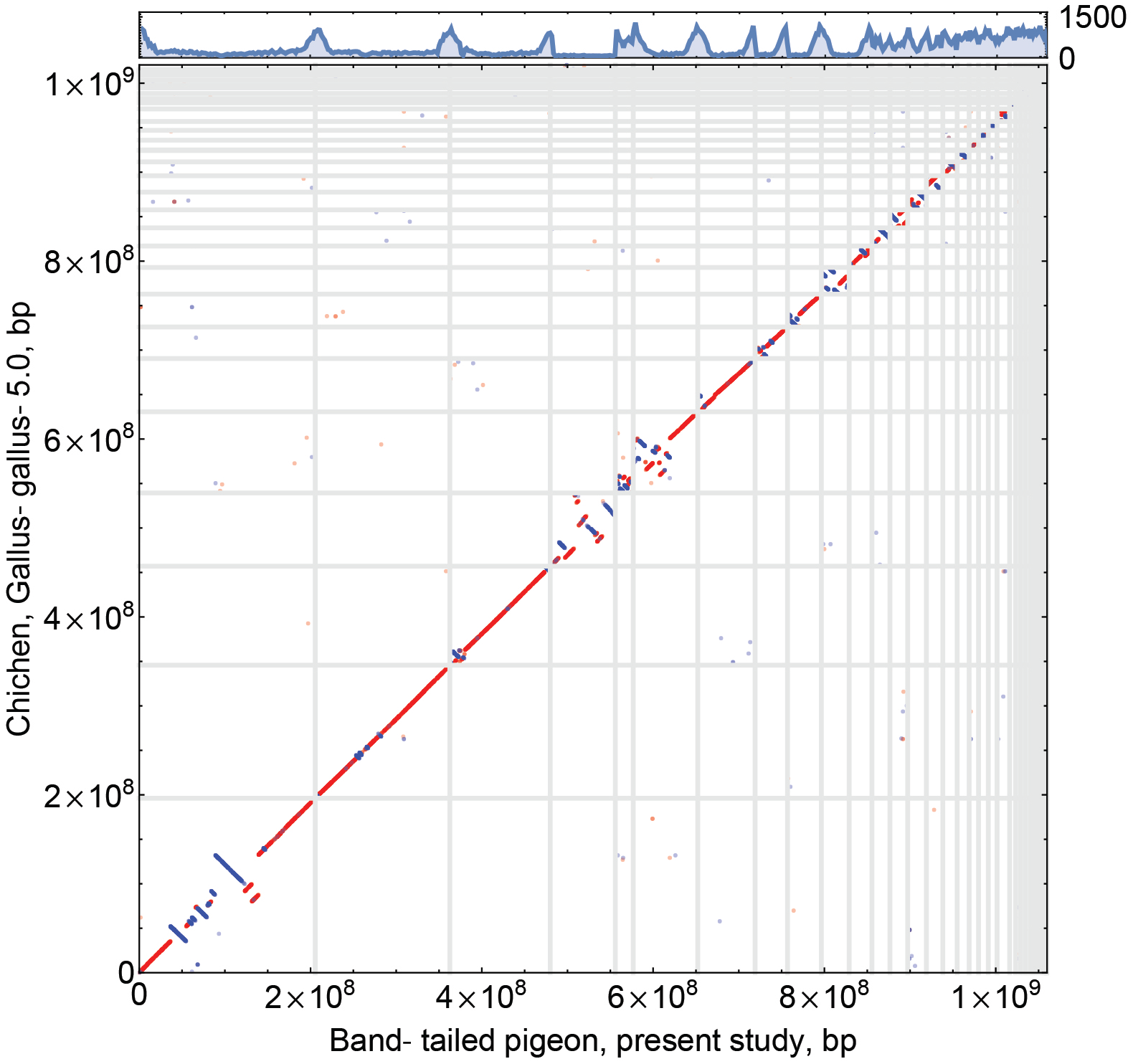
Strong conservation of synteny between the band-tailed pigeon (reassembled in this study using Juicebox Assembly Tools) and the chicken. We note very low levels of interchromosomal rearrangements (Derjusheva et al. 2004). For this analysis, the chicken (GCF_000002315.4) and the band-tailed pigeon assemblies were aligned using the LastZ alignment algorithm (Robert S. Harris 2007) using “--notransition --step=20 -nogapped” command options; the chicken assembly was used as a target. Here, we show alignment blocks with scores larger than 25,000 (Robert S. Harris 2007), with direct synteny blocks colored red, and inverted blocks colored blue. Chromosome order and orientation has been modified in order to facilitate the comparison. We also use the new assembly to revisit the question of regional variation in nucleotide diversity in passenger pigeons (Murray et al. 2017). The track on top of the synteny plot shows a total number of multiallelic sites in non-overlapping 2-Mb windows calculated from a *.vcf* file that describes the results of aligning passenger pigeon data to the draft band-tailed pigeon assembly.

**Figure S2:**
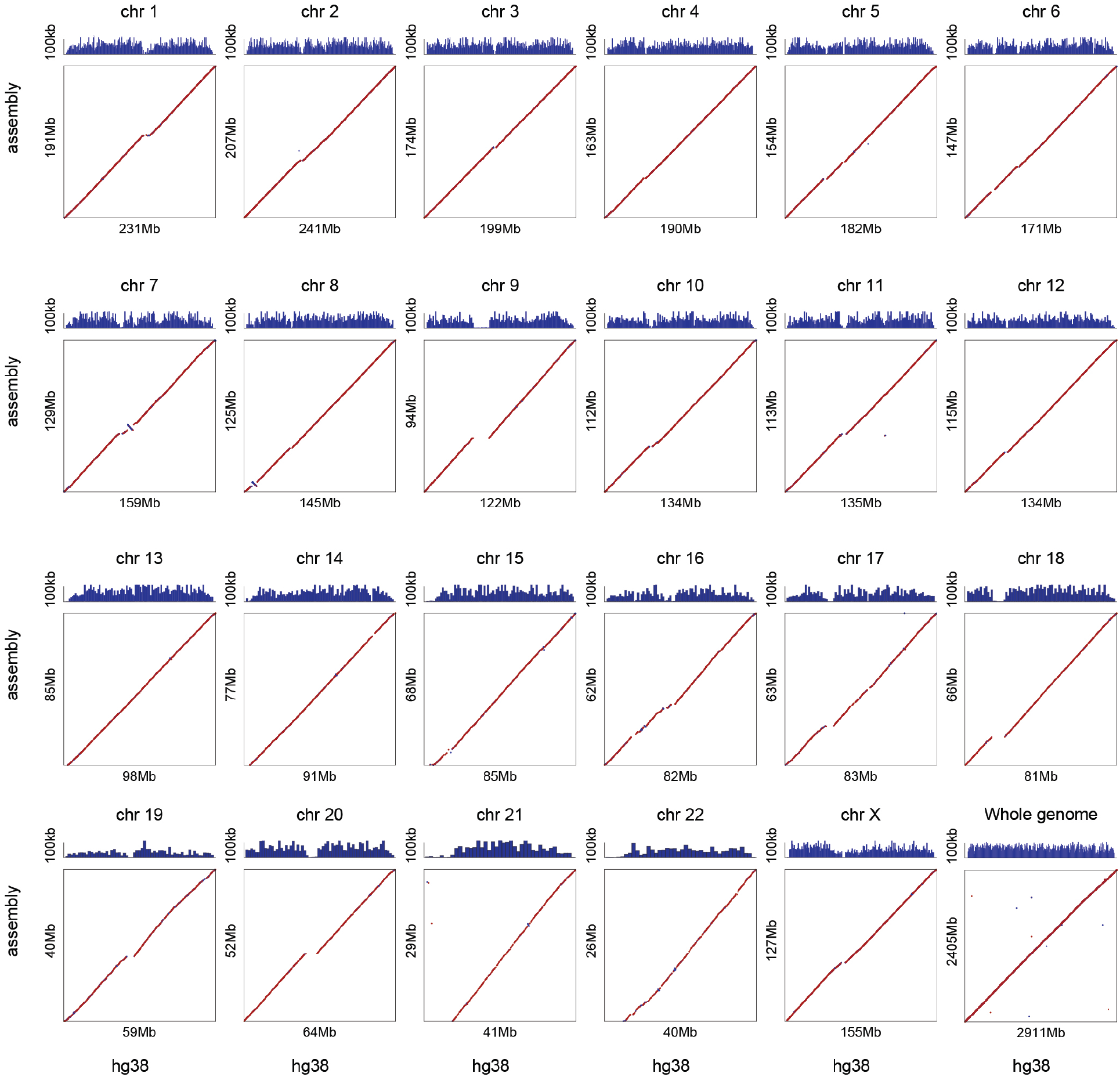
Dotplots showing alignment of chromosome-length scaffolds from hs-1k and hg38. The hg38 reference (NCBI accession number GCA_000001405.23) is shown on the X axis. The Y axis shows the 23 largest scaffolds of the hs-1k assembly; they have been ordered and oriented to match the chromosomes as defined in hg38 in order to facilitate comparison. (For the same reason, all gaps are removed in both assemblies.) Each dot represents the position of an individual resolved scaffold aligned to hg38. The color of the dots reflects the orientation of individual alignments with respect to hg38 (red indicates a match, whereas blue indicates disagreement). The track on top illustrates the scaffold N50 of the draft w2rap *de novo* assembly as a function of position (calculated in windows of 1Mb for individual chromosomes and 10Mb for the whole-genome graph). Alignment was performed using BWA (Li and Durbin 2009). The dotplots illustrate excellent correspondence between hg38 and hs-1k, with the exception of a few low-complexity regions of the human genome.

**Figure S3:**
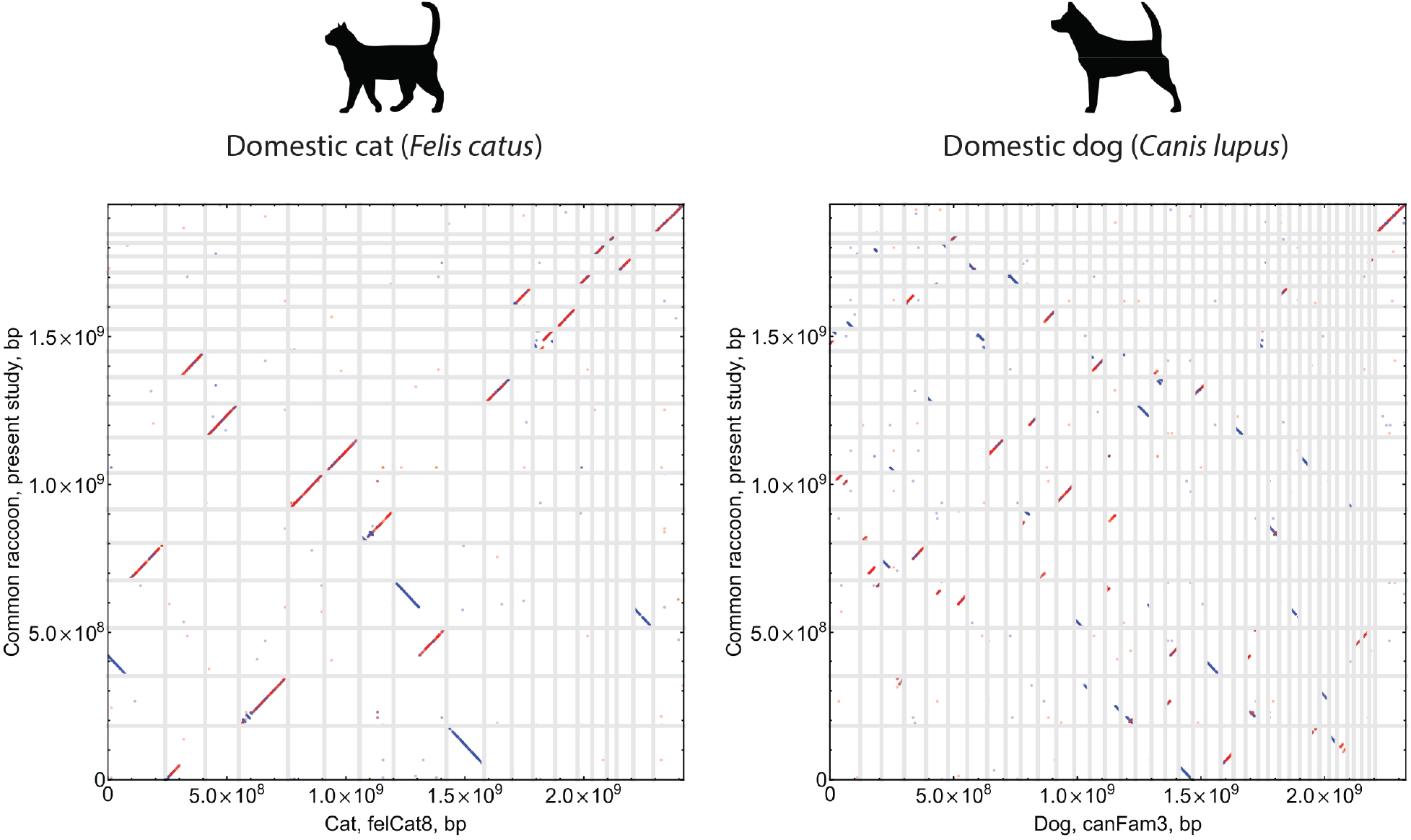
Conservation of synteny between cats, dogs, and the common raccoon (pl-1k). The analysis demonstrates a high degree of karyotype conservation between the raccoon and the cat, as well as a highly rearranged structure of the dog chromosomes. The results are in agreement with prior studies (Nie et al. 2012). For this analysis, the cat (GCF_000181335.2), the dog (GCF_000002285.3) and the common raccoon (pl-1k) genome assemblies were aligned using the LastZ alignment algorithm (Robert S. Harris 2007) using “--notransition --step=20 -nogapped” command options; the cat and dog assemblies were used as targets. Here, we show alignment blocks with scores larger than 50,000 (Robert S. Harris 2007), with direct synteny blocks colored red, and inverted blocks colored blue. Chromosome order and orientation of the common raccoon chromosomes has been modified in order to facilitate the comparison with (Nie et al. 2012).

